# Cardiac p16 Expression Following Vape Exposure with Nicotine Shows Sex-Specific Induction in Males but not Females

**DOI:** 10.1101/2024.08.01.606074

**Authors:** Abraham Grant Shain, Clarissa Savko, Sophie Rokaw, Faid Jaafar, Abigail Rieder, Morgan K. Wright, Joy A. Phillips, Nicholas Konja, Sama Michael, Haley Mathews, Gina Jerjees, Barbara Bailey, Mark A. Sussman

## Abstract

Vaping is marketed as a safe alternative to traditional cigarette smoking, but multiple studies demonstrate deleterious cardiopulmonary effects including cardiac function decline and fibrotic remodeling with alveolar size enlargement. Nicotine, a common constituent of vaping aerosol, stimulates p16 expression in pulmonary tissue but the impact on cardiac tissue remains unclear. In this study, mice were exposed to e-cigarette vape aerosol either containing nicotine (Vape Nicotine; VN) or without nicotine (Vape 0; V0). Non-exposed (No Vape; NoV) mice were used as controls. Cardiac effects were assessed by echocardiography, histology, and immunofluorescence to determine changes in function, morphology, p16, and Discoidin Domain Receptor 2 (DDR2). VN depressed cardiac function and increased collagen deposition relative to V0 and NoV. Interestingly, p16 expression was increased in cardiomyocytes and interstitial cells of male mice while remaining unchanged in females. In contrast to VN, V0 had no significant impact on cardiac function or p16 expression in males. Furthermore, collagen deposition in the V0 group was significantly lower than the VN group. Subsequent cardiac fibroblast analysis using DDR2 revealed increased expression within the V0 group relative to VN and NoV. Collectively, these findings show collagen accumulation as well as p16 expression prompted by vaping is mediated by nicotine as a constituent of vape juice. In contrast, vape aerosol alone promotes accrual of cardiac fibroblasts without concomitant changes in collagen accumulation or p16 expression. These results are the first to identify p16 induction with pathologic collagen deposition by exposure to vape aerosol containing nicotine in male cardiac tissue. The underlying basis for sex-specific differences in cardiac responses to vape aerosol exposure warrant further investigation, particularly those involving cellular and molecular changes that may lead to pathologic changes later in life.

## Introduction

The detrimental impact of traditional cigarette smoking resulted in the advent of electronic nicotine delivery systems (ENDS) such as electronic cigarettes (E-cigs) as an alternative nicotine delivery device. E-cigs vaporize e-cig juice, a mixture of propylene glycol/vegetable glycerin (PG/VG) containing flavoring and often nicotine. Additives such as children’s cereal, candy, or fruit flavoring appealing to juveniles and young adults attracted a youthful demographic to e-cigs while nicotine addiction made cessation difficult^1^. The magnitude of the problem is highlighted by self-reported high school usage of vape products representing 2.5 million American youths in 2022^2^. The health risks e-cigs pose include pathological changes to lung tissue as well as cardiac remodeling leading to diminished function^3^.

E-cig use has been linked to biological processes such as cytotoxicity, oxidative stress, inflammation, and altered gene expression in both respiratory and cardiovascular systems^4^. Nicotine inhalation modulates the cardiovascular system, including an increase in heart rate, blood pressure, and cardiac contractility via an increase in catecholamine levels. Blood flow is reduced in cutaneous and coronary vessels, promoting a hypoxic myocardial environment^5,6^. Hypoxia inhibits DNA repair, activates DNA damage response pathways, and increases reactive oxygen species (ROS)^7,8^. Genomic responses consistent with ROS activity in pulmonary tissue as a consequence of vaping were previously reported by our laboratory^9^. Additionally, chronic ROS exposure in cardiac tissue causes pathophysiological changes such as cardiac fibrosis, left ventricular hypertrophy, reduced ejection fraction, and activation of senescence pathways ultimately leading to heart failure^10–12^.

Senescence activation occurs through two cyclin-dependent kinase (CDK) inhibitor (CDKN) pathways, the CDKN2a-INK4A (p16)-retinoblastoma (Rb)-associated pathway, and the tumor-protein 53 (p53)-CDKN1a-CIP1 (p21) pathways^13,14^. Both routes lead to cell cycle arrest by inhibiting Rb phosphorylation through p16-CDK4/6 and/or p21-CDK2/4/6 binding^15–20^. Additionally, CDKN1b-KIP1 (p27) acts in the same fashion as p21^21–25^. Myocytes expressing p16 undergo hypertrophy and/or apoptosis^26,27^ while fibroblasts expressing p16 deposit increased collagen in conditions of atrial fibrillation (AF) or transverse aortic constriction (TAC), leading to impaired cardiac function^28,29^. The present study assesses pathophysiological changes that correlate the impact of vaping with nicotine on myocardial function and structure with respect to CDKN pathways.

## Materials and Methods

### Study animals

All mouse studies were performed as approved by SDSU IACUC. C57BL/6 5-wk-old male and female mice (Charles River) were housed four mice per static cage. Room temperature was held between 70-72°F and 12-hour automated light-dark cycle were constant. Rodent Maintenance Diet (Teklad Global 14% Protein) and water were provided ad libitum.

### Mouse vaping inhalation protocol

Mouse vaping protocol for this study has been previously described (Supplemental Figure 1)^9^. Vaping started at mouse age of 6-8 week (∼56 d) and continued for 9 weeks (age at conclusion equals 112–119 d). Animals were exposed for 4 h/day, 5 d/week for 9 weeks (Supplemental Figure 1A). During the adult phase of mouse life, 2.6 d are approximately equivalent to one human year. This represents the equivalent of human vaping from 16 to 37.5–40.2 year of age according to Dutta and Sengupta^30^ or the equivalent of 18–25 year of age according to Flurkey^31^. Prior to exposure mice were randomly sorted into three groups: one exposed to Peach Ice vape juice (ORGNX) with 50mg/30mL nicotine (VN), one exposed to Peach Ice vape juice (ORGNX) with no nicotine (V0), and the last exposed to room air. Peach Ice vape juice consists of 70% VG and 30% PG. Vape juice (with and without nicotine) was aerosolized by JUUL pens in whole body exposure chambers (inExpose; SCIREQ). Vaping chamber setup is represented schematically in Supplemental Figure 1A. Mice were exposed to 3 second puffs every 20 seconds at a 1.8 liters/minute intake rate. Fresh air flow delivered at 1.5 liters/minute for the 4 hour duration of the vaping profile (Supplemental Figure1B) based upon human vaping topography recommended by Farsalinos et al. concluding “4-s puffs with 20–30 s interval should be used when assessing electronic cigarette effects in laboratory experiments, provided that the equipment used does not get overheated.”^32^ In addition, puff duration parameters and frequency are within the reported range of human vaping topography from real time characterization of electronic cigarette use in the 1 Million Puffs Study^33^.

### Echocardiography

Transthoracic echocardiography images were acquired using a Vevo 2100 (VisualSonics). Chest hair was removed using Nair hair removal cream. Mice were lightly anesthetized under 3.5% isoflurane (Abbot Laboratories) using an active scavenging filtration system (Harvard Apparatus) and maintained under 1.0% isoflurane for the duration of image acquisition. Ultrasound transmission gel (Aquasonic) was used to couple the probe for imaging. Images were taken in the 2D parasternal long-axis (PSLAX) view using an MS550D transducer. Echocardiography images were analyzed using VisualSonics with SAX package. Manual heart rate measurements were taken from peak to peak of systolic ventricular movements to confirm digitally calculated heart rates. Ejection fraction (EF) and fractional shortening (FS) measurements were not taken when heartrate was below 500 or above 600 beats per minute (BPM)^34,35^.

### Tissue Preparation

Hearts were harvested for either formalin-fixed paraffin-embedded (FFPE) or flash-frozen tissues. All mice were first anesthetized with an intraperitoneal injection of ketamine. FFPE hearts were arrested in diastole with 50mM KCL and perfused with neutral buffered formalin for 10 minutes at 80 mmHg via retrograde cannulation of the abdominal aorta. Retro-perfused hearts were excised from the thoracic cavity and fixed overnight in formalin at room temperature, then processed for paraffin embedding.

### Hematoxylin & Eosin staining

Heart sections were stained as previously described^9^ to visualize morphologic and structural changes. Images were acquired with a Keyence BZ-X800 microscope. Tile scans were stitched using the Keyence software.

### Cardiac Morphology Measurements

Hematoxylin & Eosin (H&E)-stained sections imaged using Keyence analysis software were used to calculate width and area of each ventricular and the septal wall. Width was calculated by taking the average of five separate measurements evenly spaced along the vertical axis of the ventricular wall. To calculate septal and ventricular area the freehand area selection tool was used to outline each wall and then total area was graphed. Septal and ventricular length were measured by drawing a line along the endocardium and epicardium, then those two lines were averaged and graphed.

### Masson’s Trichrome staining & Collagen Deposition Analysis

For myocardial collagen deposition, the sections of heart tissue were stained following previously reported Masson’s Trichrome protocol^9^. Images were obtained by a Keyence BZ-X800 microscope. Collagen area fractions were measured using Hybrid Cell Count Analysis software within the Keyence analysis package. Only ventricular and septal tissue measurements were included in the quantitation.

### Immunofluorescence

Heart samples were formalin fixed and paraffin embedded as previously described^9,36^ Hearts were sectioned (5μm) in the coronal plane (Leica, Microtome RM2245) exposing the four-chamber view of the heart. The presence of the mitral and tricuspid valves confirmed equivalent depth of sectioning across all samples. Antibodies with working dilutions are listed in Table 1. Images were collected using a Leica SP8 confocal microscope and processed with Leica software. Stains were performed on four biological replicates per treatment group and two technical replicates. DDR2 staining colocalization with DAPI was quantified using ImageJ.

**Table 1.** Antibody Concentrations used for immunofluorescence.

| Marker | Species | Company | Catalog # | Dilution | Secondary Antibody |
| --- | --- | --- | --- | --- | --- |
| P16 | Mouse | Santa Cruz Biotech | Sc-1661 | 1:100 | DAM cy3 |
| P21 | Rabbit | Abcam | Ab188224 | 1:100 | DAM cy3 |
| P27 | Rabbit | Cell Signaling Tech | 2552 | 1:100 | DAM cy3 |
| Smooth Muscle Actin | Rabbit | Protein Tech | 14395-1-AP | 1:400 | DAR 680 |
| Smooth Muscle Actin | Mouse | Protein Tech | 67735-1-AP | 1:400 | DAM 680 |
| Vimentin | Chicken | Thermo Fisher | PAI-10003 | 1:400 | DACH 488 |
| Cardiac Troponin I | Goat | Abcam | Ab56357 | 1:400 | DAG 633 |
| DDR2 | Invitrogen | PA5-27752 | Rabbit | 1:200 | DAR cy3 |

### Statistical Analysis

Data analysis and graphical representation were performed with Prism 10 (GraphPad software) unless otherwise noted. Comparisons of treatment groups and sex differences were performed by 2-way ANOVA. Statistical significance of interactions was identified using Tukey’s post hoc multiple comparisons test.

## Results

### Cardiac functional response to vaping shows modest changes

The effect of vaping upon myocardial function was assessed by echocardiography. Long axis views of mouse hearts were taken at the completion of the vaping study prior to sacrifice and analyzed for ejection fraction (EF) and fractional shortening (FS). Representative echocardiographic images of each treatment group during systole and diastole are shown (Figure 1A). Ejection fraction between male NoV and VN trended toward significance, from 62.14% to 60.58% respectively (p<0.05). Comparisons of the remaining groups and between sexes were not significantly different (Figure 1B). Similarly, fractional shortening did not change between treatments and sexes (Figure 1C). Echocardiographic analysis indicates modest ejection fraction decline in males after VN exposure without significant loss of fractional shortening with our vaping protocol.

**Figure 1:**
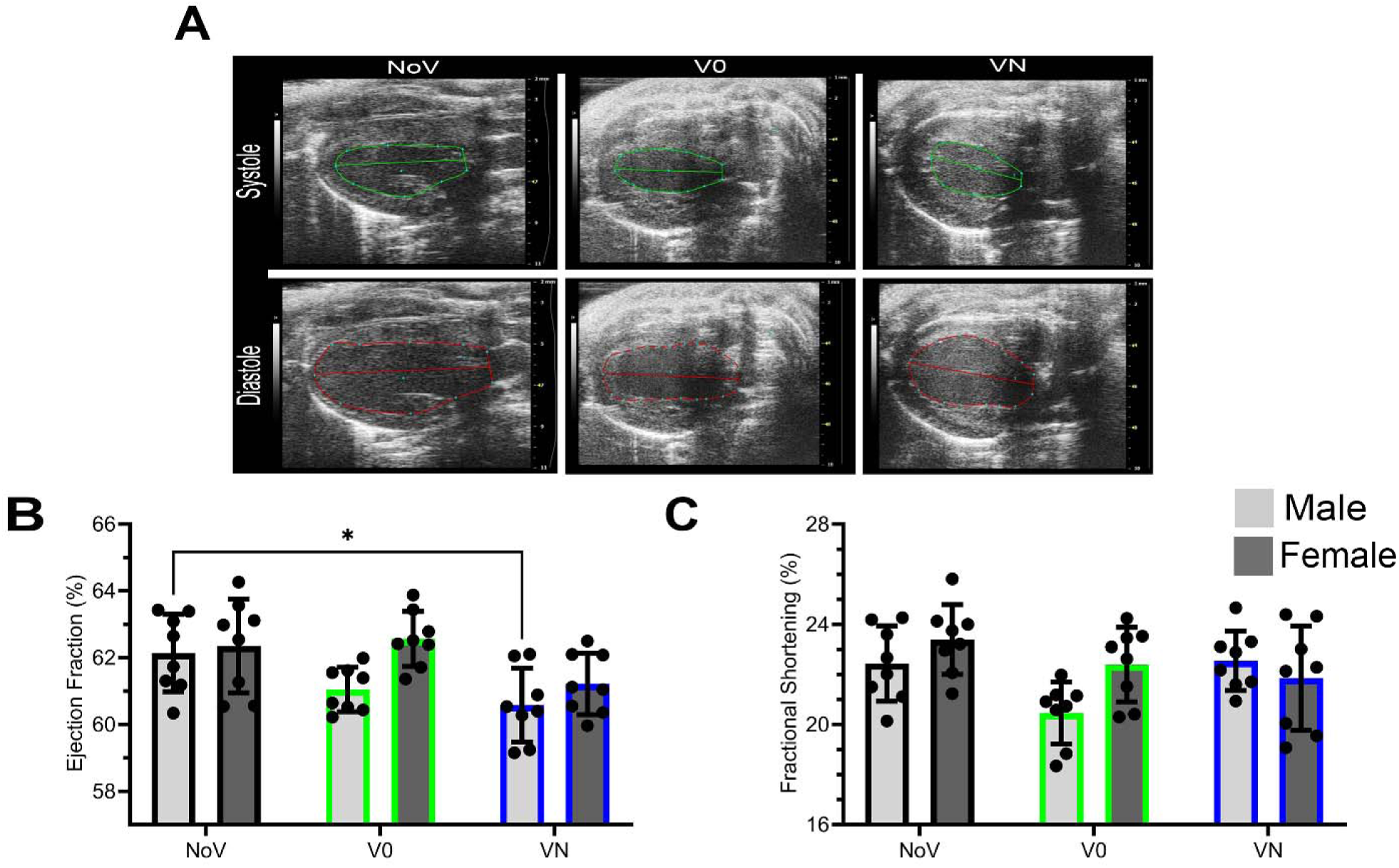
Cardiac functional response to vaping. Parasternal long axis (PSLAX) echocardiography reveals incremental changes in ejection fraction while fractional shortening remains the same. (A) Representative images of echocardiography measurements of each treatment. (B) Ejection fraction analysis. (C) Fractional shortening analysis (Male: NoV n=8, V0% n=8, VN n=8. Female: NoV n=8, V0% n=8, VN n=8). Statistical analysis performed by two-way ANOVA with Tukey’s multiple comparisons post test; each dot represents an average of three measurements per animal (*=p<0.05). No vape aerosol exposure - NoV, Vape aerosol no nicotine - V0, Vape aerosol with nicotine - VN

### Cardiac morphometric response to vaping

Echocardiographic results were further supported by subsequent *ex*-*vivo* morphometric analysis of hearts. FFPE tissue was sectioned and stained with H&E to perform morphometric analysis. Representative whole heart tile scans (Figure 2A) for all groups are shown. Measurement of left ventricular area in the V0 group decreased by 1.7 mm^2^ in males compared to females (8.1 mm^2^ versus 6.3 mm^2^ respectively; Figure 2B; p=0.002). NoV and V0 animals showed no statistically significant differences between groups or sexes (Figure 2B). Collectively, these findings demonstrate that vaping does not lead to significant cardiac remodeling relative to NoV controls.

**Figure 2:**
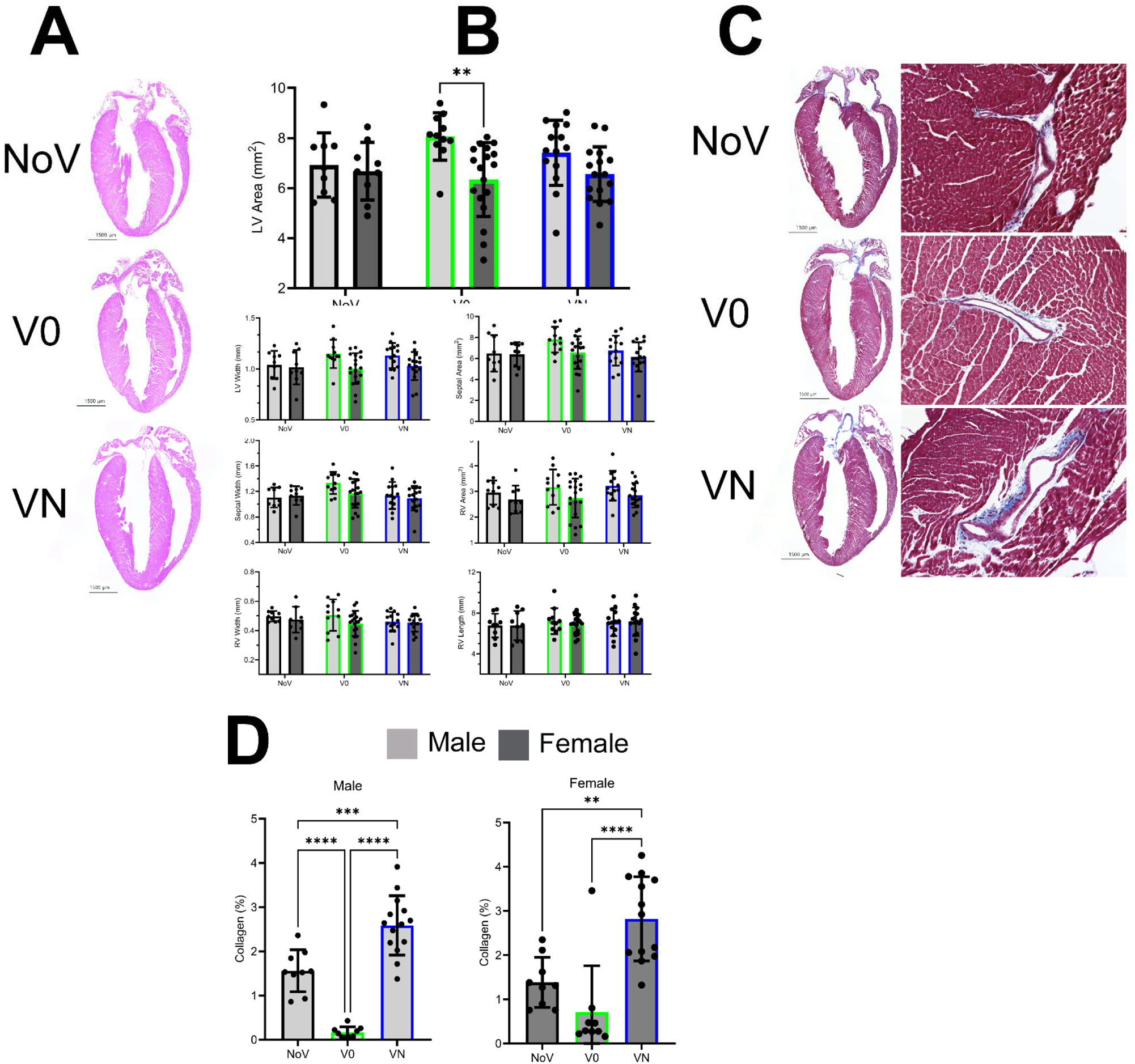
Morphometric cardiac changes after vape exposure. Hearts were stained with Hematoxylin and Eosin to measure ventricular and septal wall width and area. (A) Width and area measurements. Masson’s Trichrome used to detect collagen deposition in the septal and ventricular walls. (B) Representative images. (C) Representative tile scans with magnified images of collagen deposition surrounding blood vessels. (D) Collagen deposition per total area of ventricle and septum. (Male: NoV n=9, V0 n=11 VN n=14. Female NoV n=9, V0 n=18, VN n=16). Statistical analysis = One-way ANOVA with Tukey’s multiple comparisons post test (* P<0.05, **P<0.01, ***P<0.001, ****P≤0.0001). No vape aerosol exposure (NoV), Vape aerosol no nicotine (V0), Vape aerosol with nicotine (VN).

### Collagen accumulation promoted by nicotine

Collagen deposition, a hallmark of cardiac remodeling, was assessed using Masson’s Trichrome stain. Representative tile scans and corresponding high magnification images are shown (Figure 2C). VN heart sections show increased collagen staining relative to NoV and V0 which were comparable (Figure 2D). Average collagen levels of VN significantly increased for both sexes relative to NoV from 1.6% to 2.6% in males and 1.4% to 2.8% in females (Male p<0.0.01, Female p<0.01). V0 males show a significant decrease in collagen deposition compared to both NoV (p=0.0001) and VN (p<0.0001) exposed mice. Collagen deposition in VN females was significantly increased relative to V0 females (p<0.0001). No sex-specific differences when comparing each treatment group were observed across all three treatments. Overall, these findings are consistent with nicotine exposure driving collagen fibrosis in VN treatment group relative to NoV and V0 mice.

### p16 expression increased by nicotine in males but not females

Senescence markers p16, p21, and p27 were assessed via immunofluorescence confocal microscopy to identify cellular distribution. Paraffin embedded hearts were sectioned and labeled with antibodies to label myocytes, interstitial cells, senescence marker, and a nuclear stain for quantitative analysis using 8 20x left ventricular free wall images. Representative images show nuclear p16 localization associated with either myocytes or interstitial cells (Figure 3A). Cardiomyocytes of VN males showed significantly higher (p<0.0001) frequency of p16^+^ (5.16 per field) compared to VN females (0.34 per field), NoV males (1.72 per field), or V0 males (0.06 per field). Similarly, interstitial cells of VN males showed a significant increase of p16^+^ (7.8 per field) compared to VN females (1.8; p<0.0001), V0 males (0.13 per image; p<0.0001) or NoV males (5.06; p<0.001) (Figure 3B). Furthermore, sex comparison in the NoV groups showed significantly higher p16^+^ interstitial cells in males relative to females (5.06 to 2.41, respectively; p=0.0014). In contrast to males, females exhibited no significant differences in average p16^+^ between treatments or groups for either cardiomyocytes or interstitial cells. In summary, nicotine promotes p16^+^ in male hearts for both cardiomyocytes and interstitial cells. Additional senescence markers including p21 and p27 were not increased after VN exposure (data not shown). Therefore, nicotine promotes p16 expression in male but not female hearts.

**Figure 3:**
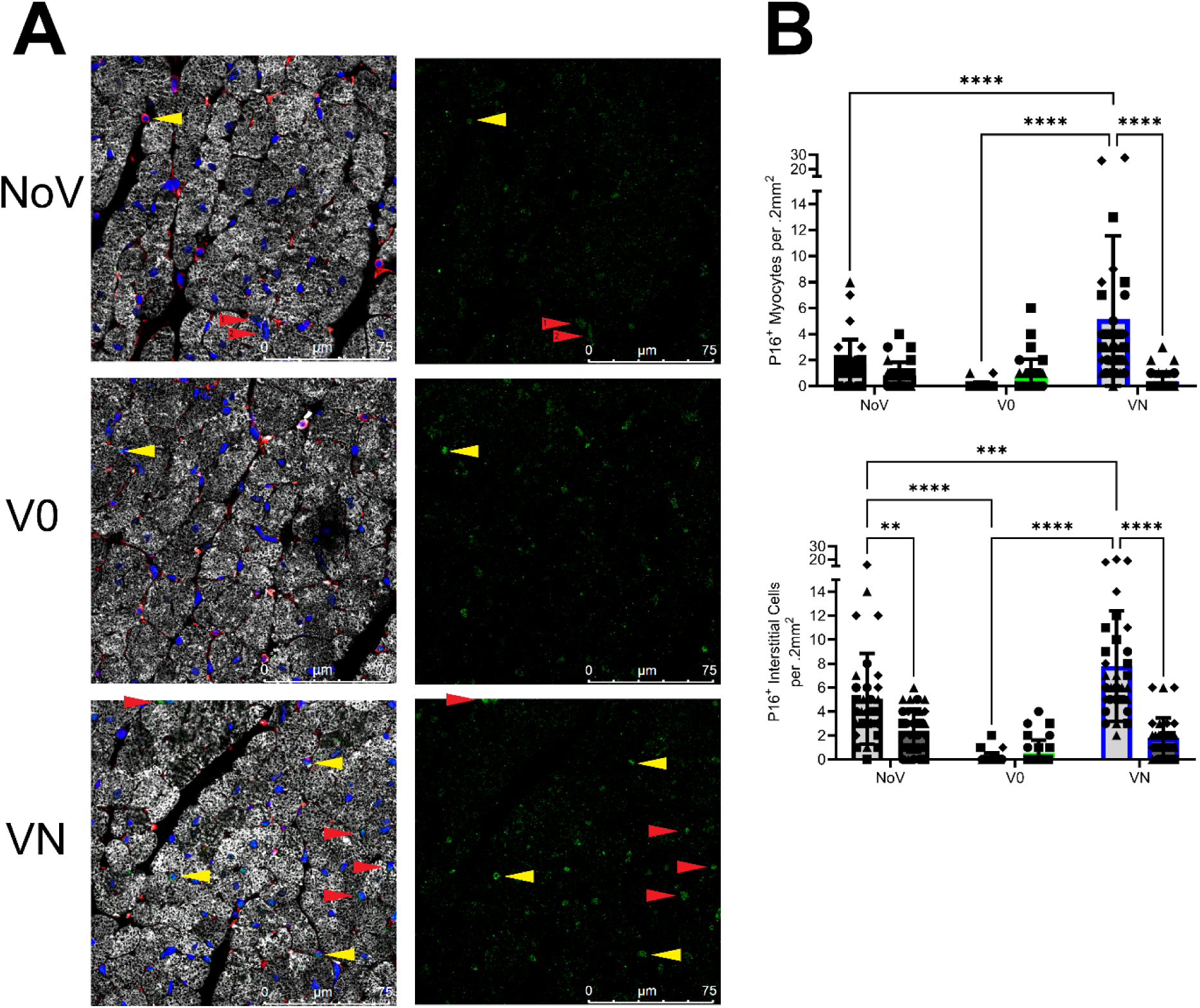
Nicotine promotes p16^+^ immunoreactivity in male but not female hearts. Immunofluorescence (IF) labeled heart sections for p16^+^ (green) in cardiomyocytes (cardiac troponin I; white) or interstitial cells (vimentin; red) colocalized with nuclei (blue). Cardiomyocyte nuclei colocalized with p16^+^ (red arrows). Interstitial cell nuclei colocalized with p16 (yellow arrows). All representative images are taken at 63x magnification. Cell count graphs represent eight separate 20x magnification images per heart, the number of total counted cells was averaged per image (0.2mm^2^). (A) Representative IF images, merge on the left and senescence marker single channel on the right. (B) p16 myocyte and interstitial count. Various symbols in the graph refer to individual hearts in the analysis. Two-way ANOVA with Tukey’s multiple comparisons test (**p<0.01, ***p<0.001, ****p<0.0001; n=4). No vape aerosol exposure (NoV), Vape aerosol no nicotine (V0), Vape aerosol with nicotine (VN).

### DDR2 colocalization with DAPI in cardiac tissue

DDR2 expression was assessed with immunofluorescent confocal microscopy together with DAPI colocalization to label nuclei. Using paraffin sections from the same hearts as the p16 distribution analysis cells were labeled with myocyte, interstitial (DDR2 and Vimentin), and DAPI. Representative images show DDR2 labeling is brightest in the V0 group and virtually absent in NoV and VN exposed hearts. In V0 cardiac tissue DDR2 localizes to nuclei and is observed in both male and female mice (Figure 4A). Quantitative analysis was performed using 12 40x left ventricular free wall images immunolabeled with cTnI (myocytes), DDR2 (interstitial cells), and DAPI (nuclei). Percentage overlap between DDR2 and DAPI was calculated using an ImageJ macro processor. Significant increases in DDR2/DAPI overlap in the male and female V0 group relative to both NoV and VN groups were present (p<0.0001). Female VN left ventricular tissue sections also had less frequent DDR2/DAPI overlap than NoV (p<0.001). This data indicates vape aerosol triggers DDR2 expression, however, vape with nicotine suppresses DDR2 expression to similar levels of non-vape exposed animals.

**Figure 4:**
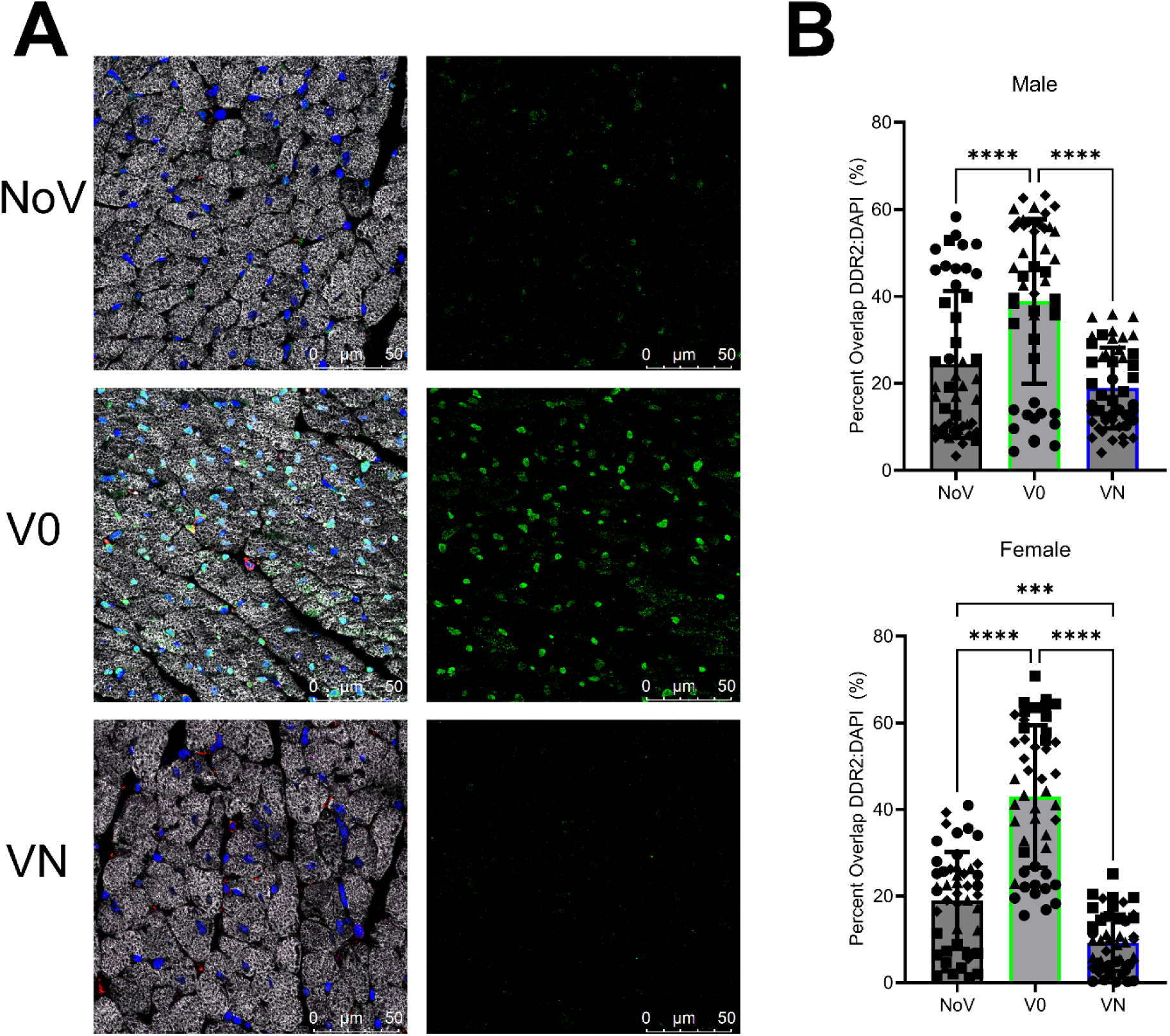
DDR2 expression increases after exposure to vape aerosol without nicotine. Percent overlap graph represents summation of 12 40x magnification images per animal, 4 animals per group. (A) Representative IF labeled heart sections of DDR2 immunolabeling with DAPI taken at 63x magnification. (B) Percent overlap of DDR2 and DAPI expressed as the percentage of DDR2 fluorescence overlapping the total DAPI fluorescence. Individual animals in the analysis are represented by differing symbols in the graph. Two-way ANOVA with Tukeys multiple comparisons test (***p<0.001, ****p<0.0001; n=4). No vape aerosol exposure - NoV, Vape aerosol no nicotine - V0, Vape aerosol with nicotine – VN

## Discussion

The meteoric rise in vaping has been particularly alarming due to adoption by individuals (the “never smokers”) with no prior history of cigarettes that include a youthful demographic attracted by targeted advertising and the lure of experimentation^37–41^. Conflicting messages from pro-vaping versus anti-vaping advocacy groups coupled with disdain or mistrust of scientific literature lead to indifference or skepticism regarding health-related concerns^42,43^. The addictive nature of nicotine contained in cigarettes and most vape juice products leads to long term use that, rather than encouraging cessation, can escalate into greater dependency and dual use^44^. While harm reduction through cessation of combustible cigarette smoking improves public health, there is abundant evidence that encouraging vaping as an alternative has created new health risk concerns and expanded the reach of Big Tobacco products^45–47^.

Vaping provokes sex-specific depression of cardiac function as well as activating oxidative stress, inflammation, and fibrosis^48–50^. Consistent with these effects we found nicotine-dependent fibrosis with collagen deposition significantly increased in VN exposed animals relative to NoV and V0 (Fig 2C,D). Although nicotine in tobacco smoke increases collagen deposition^50^ there is evidence of decreased accumulation from e-cig usage^51,52^. Vaping with nicotine resulting in increased collagen deposition has not been previously published for the myocardium. Importantly, cardiac structure was not significantly impacted in V0 or VN groups of the present study yet vaping with nicotine showed a slight decline in cardiac function in males only (Figure 1A, Figure 2A,B). These findings are consistent with early detrimental changes prompted by nicotine that may impair cardiac function as well as capacity to recover from injury later in life. The long-term consequences of such changes remain to be determined.

Nicotine from e-cigs induces DNA damage and impairs DNA repair mechanisms promoting oxidative stress which drives senescence^53–57^. VN males exhibited more p16^+^ interstitial cells after VN exposure compared to V0 males and VN females. Additionally, males exposed to VN had significantly more p16^+^ cardiomyocytes than their female or V0 male counterparts (Figure 3A,B). Consequences of p16 cardiomyocyte expression remain unclear with potential for hypertrophic remodeling^58^ as well as inhibition of hypertrophy^59^. Alternatively, p16 expression has been associated with cardiomyocyte apoptosis which erodes cardiac structure and function through multiple mechanisms including blunted DNA synthesis together with diminished myocyte shortening, contractile velocity, and Ca^2+^ reuptake reminiscent of cardiac aging^60^. Aged cardiomyocytes expressing p16 suffer from impaired resistance to stress due to collective effects of reduced and altered mitochondrial respiratory chain leading to ROS accumulation. Chronic ROS activity compromises healthy cardiomyocytes by propagating senescence and activation of p16^10,61,62^. Therefore, this study may show early accumulation of cardiomyocyte senescence in male hearts but stops premature of inevitable tissue decline with natural aging that could be accelerated by vaping with nicotine.

Interstitial cell, notably fibroblast, senescence is activated by transient p21 expression, temporary p27 upregulation, then long term p16 expression^63^. Senescent fibroblasts deposit more collagen stiffening ventricular tissue and impairing ventricular wall movement^61,63,64^. Fibroblasts also express nicotinic alpha 7 receptor (α7nAChR), the receptor which mediates nicotine binding. α7nAChR activates noradrenaline secretion stimulating β-adrenergic receptors (β-AR) which mediates collagen deposition^65–67^.

DDR2 is a receptor tyrosine kinase expressed on resident cardiac fibroblasts and interacts with collagen to facilitate tissue remodeling. DDR2’s canonical function involves promoting growth, migration, and differentiation. Resident fibroblasts distributed interstitially throughout myocardial tissue maintain tissue structure by depositing collagen I, III, V, vimentin, and fibronectin^68^. Reports demonstrate both proliferative and antiproliferative actions of DDR2^69–76^, however only one investigates DDR2’s interaction with cell cycle inhibitors^77^. To our knowledge no study to date has investigated how vaping affects DDR2 expression in cardiac tissue. In the present study, exposure to vape aerosol without nicotine (V0 group) promoted DDR2 expression but did not correspond to collagen deposition. Together this data implies the cellular response to vape aerosol without nicotine provokes DDR2 expression but does not impact collagen production. In comparison, addition of nicotine to the vape aerosol suppresses the increase of DDR2 but activates collagen production and fibrosis. DDR2 upregulation activate cell cycle through CDK4/5 and Rb^77^, so DDR2 increases in the V0 group may operate through a similar mechanism to increase interstitial cell numbers without promotion of collagen deposition by nicotine-mediated fibroblast activation.

Cardiac fibroblast-mediated collagen deposition is increased following adrenergic drive^78^. Although males of the VN group possess more p16 positive interstitial cells than similarly exposed VN females there were no evident fibrotic differences between sexes (Figure 2D). Fibrotic development is a long-term chronic process that may not have had sufficient time to evolve in our short-term study. Of particular interest and concern is how and whether these early changes following vaping with nicotine could contribute to accelerated cardiac decline over lifespan as is the case for conventional cigarette smoking^79–81^. Cardioprotective pathways in females could account for lower level of p16^+^ relative to males in the VN groups.^82,83–86^.

This study establishes elevated male p16 accumulation in both cardiomyocytes and interstitial cells after exposure to vape aerosol containing nicotine. p16-mediated cell cycle arrest would be an ideal starting point to identify how nicotine lowers DDR2 expression in hearts of nicotine-exposed males. p16 binds to CDK4/6 inhibiting phosphorylation of pRb, pushing pRb to a hypophosphorylated state sequestering E2F transcription factors ultimately arresting the cell cycle in the G1 phase^87^. Corroboration of immunofluorescence results with isolated enriched cardiomyocyte or interstitial cells preparations could be performed to measure p16 or DDR2 levels in future studies.

Chronic impact of VN exposure upon the heart are deleterious based upon our findings, although long term effects of vaping with nicotine remain poorly understood. Mice of advanced age exposed over an extended time span would need to be compared with age-matched subjects vaped using our protocol to assess the longterm effects. This study found that moderate vaping with or without nicotine does not severely affect cardiac function in young to middle aged mice. However, nicotine exposure drives early cardiac fibrosis regardless of sex. Nicotine also increases male cardiac p16 accumulation indicating this process is independent of fibrotic deposition. The observation of DDR2 accumulation with a lack of collagen deposition in V0 warrants further study to understand the interplay between vape aerosol exposure versus the role of nicotine in pathogenesis. Therefore, this study serves a dual purpose identifying the novel finding of nicotine induced fibrosis with the added feature of sex-specific p16+ cells accrual in males but not females. Vaping may pose as yet unresolved cardiac risk particularly for males relative to females which may arise later in life.

## Non-standard Abbreviations and Acronyms

e-cig: Electronic Cigarette
p16: CDKN2a-INK4a-retinoblastoma-associated protein
p21: CDKN1a-KIP1
p27: CDKN1b-KIP1
DDR2: Discoidin Domain Receptor 2
PG/VG: Propylene glycol/vegetable glycerin
V0: E-cigarette aerosol without nicotine
VN: E-cigarette aerosol with nicotine
NoV: Non-e-cigarette exposed

## Acknowledgements

MA Sussman is a recipient of funding from the California Tobacco Related Disease Research Program (Pilot award T31IP1790 and Impact award T31IR1585). The author extends his deepest appreciation to SDSU for providing University Grant Fellowship funding, and members of the Sussman Laboratory who provide invaluable assistance and expertise in developing vaping-related studies and information.

---

**Supplemental Figure 1:**
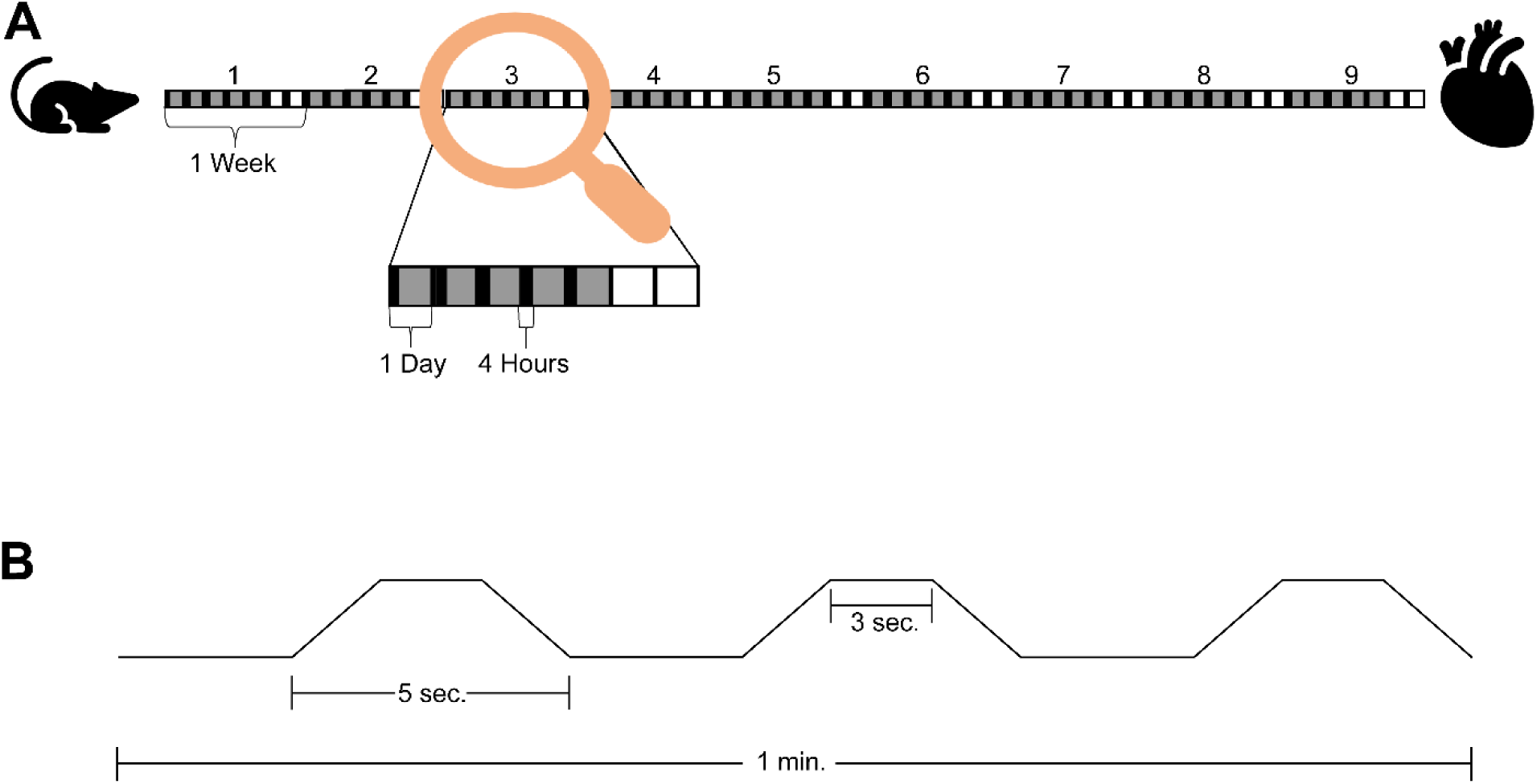
Time course and puff map of vaping study. (A) Time course of vaping study. Grey boxes indicate days mice were vaped and white boxes indicate days they were not. Black boxes indicate hours within the day mice were vaped. The time course lasted 9 weeks ending with echocardiography analysis followed by heart dissection. (B) Puff topography of 1 minute of vaping. A total of three puffs per minute were given, with 5 second puffs and 15 seconds of rest. Each puff requires 1 second of pump buildup, 3 seconds of aerosolizing, and one second of pump decline.

## Notes

### Competing Interest Statement

The authors have declared no competing interest.

### Summary of Updates

An acknowledgements section was added to this manuscript.

